# QUBIC2: A novel biclustering algorithm for large-scale bulk RNA-sequencing and single-cell RNA-sequencing data analysis

**DOI:** 10.1101/409961

**Authors:** Juan Xie, Anjun Ma, Yu Zhang, Bingqiang Liu, Changlin Wan, Sha Cao, Chi Zhang, Qin Ma

**Author notes:** To whom correspondence should be addressed. Correspondence is also addressed to Chi Zhang.

## Abstract

The combination of biclustering and large-scale gene expression data holds a promising potential for inference of the condition specific functional pathways/networks. However, existing biclustering tools do not have satisfied performance on high-resolution RNA-sequencing (RNA-Seq) data, majorly due to the lack of (*i*) a consideration of high sparsity of RNA-Seq data, e.g., the massive zeros or lowly expressed genes in the data, especially for single-cell RNA-Seq (scRNA-Seq) data, and (*ii*) an understanding of the underlying transcriptional regulation signals of the observed gene expression values. Here we presented a novel biclustering algorithm namely QUBIC2, for the analysis of large-scale bulk RNA-Seq and scRNA-Seq data. Key novelties of the algorithm include (*i*) used a truncated model to handle the unreliable quantification of genes with low or moderate expression, (*ii*) adopted the mixture Gaussian distribution and an information-divergency objective function to capture shared transcriptional regulation signals among a set of genes, (*iii*) utilized a Core-Dual strategy to identify biclusters and optimize relevant parameters, and (*iv*) developed a size-based *P*-value framework to evaluate the statistical significances of all the identified biclusters. Our method validation on comprehensive data sets of bulk and single cell RNA-seq data suggests that QUBIC2 had superior performance in functional modules detection and cell type classification compared with the other five widely-used biclustering tools. In addition, the applications of temporal and spatial data demonstrated that QUBIC2 can derive meaningful biological information from scRNA-Seq data. The source code for QUBIC2 can be freely accessed at https://github.com/maqin2001/qubic2.

## INTRODUCTION

As next-generation sequencing technologies have become more affordable in these years (1,2), it is possible to generate large-scale biological data with higher resolution, better accuracy, and lower technical variation than the array-based counterparts (3,4). RNA-Seq measures the abundance of RNA transcripts, giving rise to genome-scale gene expression data in a biological sample at a given moment (5). Nowadays, researchers can isolate individual cells from complex organisms and measure transcriptional activity using single-cell sequencing. Hundreds of RNA-seq data sets with more than hundreds of sample were emerged in the public domain in the past five years, and their tremendous values have been confirmed in many research areas, e.g., elucidation of cell-type-specific gene regulatory networks (6) and cancer & complex diseases (7-9).

The abundance of gene expression datasets provides an opportunity to computationally identify condition based functional gene modules (FGMs), each of which is defined by a similar expression patterns over a certain gene set, which tend to be functionally related or co-regulated by the same transcriptional regulatory signals (TRSs) under a specific condition. Thus, successfully derivation of the FGMs may grant a higher-level interpretation of gene expression data, improve functional annotation of genes, facilitate inference of gene regulatory relationships, and provide a better mechanism level understanding of diseases such as cancer. The identification of FGMs can be naturally modeled as a specific data pattern over unknown subset of genes and samples, and solved with a bi-clustering approach (10), a two-dimensional data mining technique that can simultaneously identify co-expressed genes under a subset of conditions (i.e., samples or cells). This unique feature makes it more useful than clustering when applied to large-scale gene expression data, as genes are usually co-expressed under certain instead of all conditions.

Besides the identification of FGMs in bulk tissue data, a similar formulation may also be applied to scRNA-Seq data, to identify individual cells or cell types as well as their complex interactions under specific stimuli, e.g., cell types classification and clustering. In multicellular organisms, biological function emerges when heterogeneous cell types form complex organs (11). Investigations into organ development, cell function, and disease microenvironment highly depend upon an accurate identification and categorization of cell types, sometimes along with their temporal and spatial features (12). Traditionally, a cell type was predicted based on morphological properties or marker proteins, yet this method failed to characterize the full diversity of cells. scRNA-Seq data provides the possibility to group cells based on their genome-wide transcriptome profiles, and several studies have already been carried out using scRNA-Seq data to identify novel cell types, proving its power to unravel the full diversity of cells in human and mouse (13). Mathematically, the problem of scRNA-seq based cell types classification can be naturally formulated as biclustering problems, since the essence is to find sub-populations of cells sharing common expression patterns among subsets of genes.

Substantial efforts have been made in biclustering algorithm and tool development since 2000 (14-26), and a few review studies have provided considerable guidance in choosing suitable algorithms in different contexts (27-29): Eren *et al.* compared 12 algorithms and concluded that our previously developed method, QUBIC, is one of the top performed methods, as it has achieved the highest performance in synthetic datasets and captured a high proportion of enriched biclusters on real datasets, comparing to Plaid, FABIA, ISA and Bimax, which were also recommended for capturing upregulated biclusters (27). In 2018, Saelens *et al*. ranked ISA, FABIA and QUBIC as the top biclustering methods in terms of predicting gene modules from human and/or synthetic data (30).

Although numerous bi-clustering methods have been developed for gene expression data analysis, the most existing algorithm are designed and evaluated using microarray rather than RNA-Seq data. One of the unique features of gene expression data derived from RNA-Seq, especially the scRNA-Seq data, is the massive zeros (up to 60% of all the genes in a cell have read counts being zeros) (31,32). The normalized read counts roughly follow lognormal distributions; however, the raw zero counts of specific genes will lead to negative infinity after logarithmic transformation (33-36), resulting in unquantifiable errors. Therefore, the biclustering methods that are successful for microarray cannot be directly applicable to RNA-Seq data (37), and novel methods taking full consideration of characteristics of RNA-Seq data are urgently needed in the public domain.

In this paper, we developed a novel bi-clustering algorithm, namely QUBIC2, for large-scale RNA-seq data analysis. We demonstrated the performance of QUBIC2 on capturing FGMs by applying it to four datasets and benchmarking against five widely used biclustering algorithms. QUBIC2 turned out to be a superior player as it has identified a significant higher proportion of enriched and diverse FGMs. Besides, QUBIC2 can also identify cell types with a higher accuracy, comparing to the five biclustering tools and SC3, a state-of-the-art cell clustering method. Furthermore, we also illustrated the application power of QUBIC2 on inferring time- and spatial-related insights from two temporal and two spatial scRNA-seq datasets.

## RESULTS

### Overall design of QUBIC2

Inheriting the qualitative representation and graph-theory based model from QUBIC (19), QUBIC2 has four unique features: (*i*) developed a rigorous truncated model to handle the unquantifiable errors caused by zeros, and used a reliable qualitative representation of gene expression to reflect expression states corresponding to various TRSs; (*ii*) integrated an information-divergence objective function in the biclustering framework in support of functional gene modules identification; (*iii*) employed a Core-Dual strategy to optimize consistency level of a to-be-identified bicluster; and (*iv*) developed a robust *P*-value framework to support statistical evaluation of all the identified biclusters. Details of these four features are showcased as follows (Figure 1).

**Figure 1.**
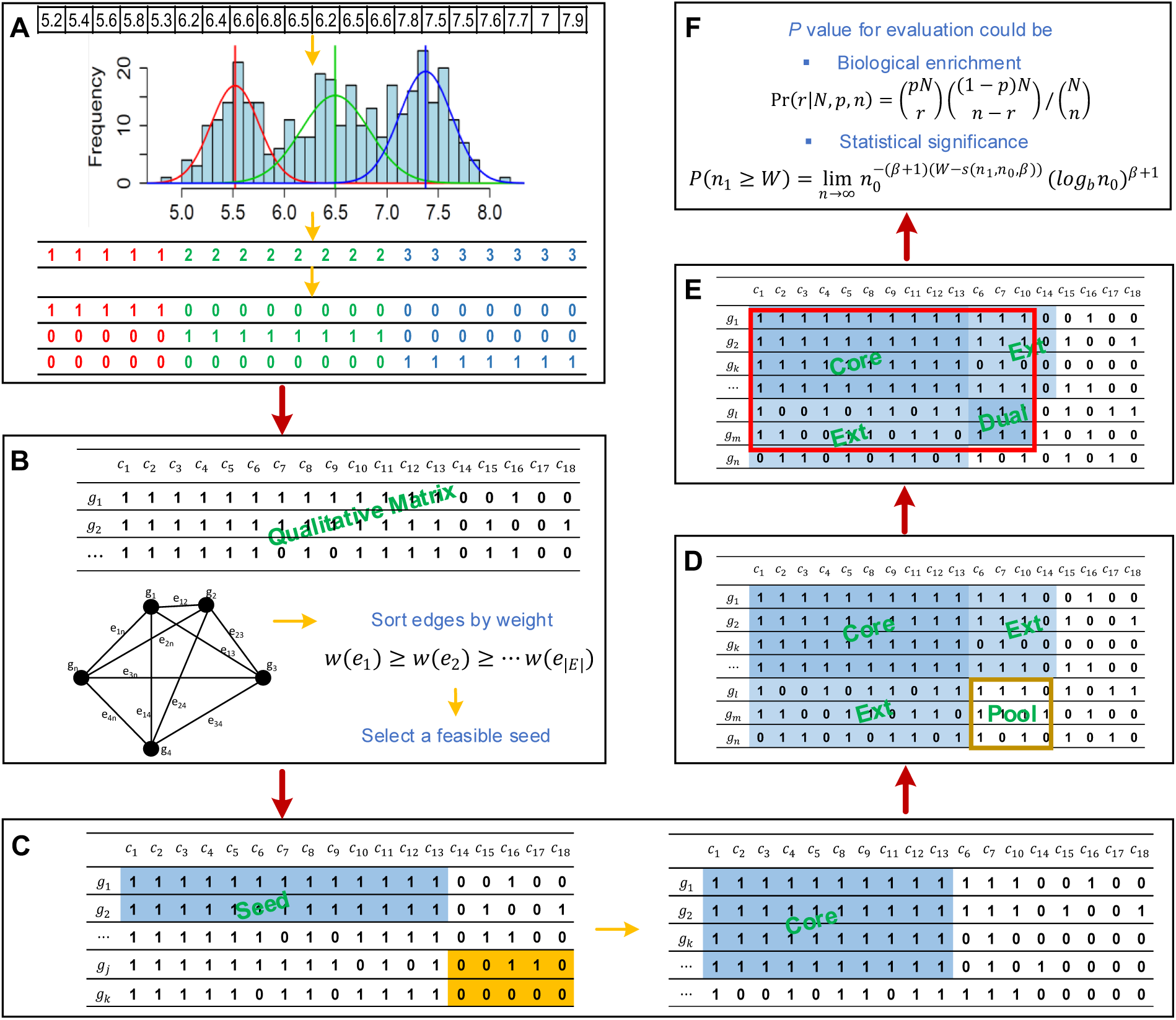
QUBIC2 workflow. **A**. Discretization of gene expression data from RNA-Seq. The LTMG model will be applied to fit each gene’s expression profile. A representing row for each gene will be generated with integers denote the most likely component distribution that each value belongs to. Then this representing row will be split into multiple rows. Finally, a binary representing matrix will be generated; **B**. Graph construction and seed selection. A weighted group will be constructed based on the representing matrix from A. By sorting the edges in decreasing order of their weight, and an initial seed list will be obtained. QUBIC2 will select a feasible seed from the list; **C**. Build an initial core based on the selected seed. During seed expand, QUBIC2 will search for genes with higher weight with the seed. In case of two genes have the same weight, the one with higher KL score will be selected. Thus, gene k (KL=0.1914) instead of gene j (KL=0.0622) will be added to the core first; **D**. Expand core and determine pool. QUBIC2 will expand the core vertically and horizontally to recruit more genes and conditions under a preset consistency level, respectively. The intersected zone created by extended genes and conditions as a Dual searching pool (brown box); **E**. Dual search in the pool and output the bicluster with genes and conditions that come from Core and Dual as final bicluster (red box); **F**. Statistical evaluation of identified biclusters based on either biological annotations or the size of the bicluster.

A mixture of left-truncated Gaussian distributions (LTMG) model was designed to fit the RNA-Seq data, rather than discarding zeros or adding a small constant to original counts (34,38). The basic idea is to treat the large number of observed zeros and low expressions as left censored data in the mixture Gaussian model of each gene (39,40), assuming that the observed frequency of expressions on the left of the censoring point should be equal to the area of the cumulative distribution function of the mixture Gaussian distribution left of the censoring point. Furthermore, we assumed that a gene should receive *K* possible TRSs under all the conditions, and its expression profile would follow a mixture of *K* left truncated Gaussian distributions. The LTMG model was applied to fit the expression value of each gene and the gene expression value under a specific condition was labeled to the most likely distribution. Accordingly, a row consisting of discrete values (1,2, ⋯, *K*) for each gene was generated (Figure 1A). Then this qualitative row was split into *K* new rows, such that in the *i*^th^ row those labeled initially as *i* are labeled as 1, while the rest were labeled as 0. Finally, a binary representing matrix *M*_*R*_ was generated.

A weighted graph *G* = (*V, E*) was constructed based on *M*_*R*_, where nodes *V* correspond to genes, edges *E* connecting every pair of genes (Figure 1B). The edge weight indicates the similarity between the two corresponding genes, which is defined as the number of conditions in which the two genes have 1s in *MR*. Intuitively, two genes from a bicluster should have a heavy edge in *G* innately while two random genes may have a heavy edge only accidentally. Hence, a bicluster should correspond to a maximal subgraph of *G*, with edges typically heavier than the edges of an arbitrary subgraph. Identifying all the biclusters equals to identifying all the heavy subgraphs in *G*, which is an NP-hard problem. Therefore, a heuristic strategy was designed as follows.

The algorithm would iterate a seed list (*S*), which is the sorted list of edges in *G* in the decreasing order of their weights (i.e., *w*(*e*_1_) ≥ *w*(*e*_2_) ≥ ⋯, *w*(*e*_|*E*|_)). An edge *e*_*ij*_ = *g*_*i*_*g*_*j*_ is selected as a seed if and only if at least one of *g*_*i*_ and *g*_*j*_ is not in any previously identified biclusters, or *g*_*i*_ and *g*_*j*_ are in two nonintersecting biclusters in terms of genes. QUBIC2 first built a core bicluster from a seed and then expanded to recruit more genes and conditions into a to-be-identified bicluster, until the Kullback-Leibler divergence score (KL score) was locally optimized. It was proposed based on the assumption that the difference between a bicluster and its background should be larger than the difference between an arbitrary same-size submatrix and its background. The KL score of a bicluster was designed to quantify this difference as the larger of the difference was, the larger of the score is (Figure 1C. See **Methods** for details).

During bicluster expansion, the algorithm controlled the consistency level for a bicluster, which is defined as the minimum ratio of the number of 1s in a column/row and the number of rows/columns in the bicluster. In QUBIC, a pre-specified value c (0<c≤1.0) was used to control the overall consistency level of the bicluster. While this parameter was dynamically optimized by a Dual searching method in QUBIC2 (Figure 1D-E), giving rise to a submatrix (*I, J*) of *M*_*R*_ (i.e., a bicluster) with optimized consistency level and maximal KL score can be identified. Biclusters expanded using Dual strategy tend to be more significant than those without Dual (See Example S1 in Supplementary File 1).

Furthermore, for the first time, a statistical framework based on the size of the biclusters was implemented to calculate a *P*-value for each of the identified biclusters. The problem of assessing the significance of identified biclusters was formulated as calculating the probability of finding at least one submatrix enriched by 1 from a binary matrix with given size, with a beta distribution employed during the process. This *P*-value framework enables users systematically evaluate the statistical significance of all the identified biclusters, especially for those from less-annotated organisms (Figure 1F).

### Functional gene modules detection from RNA-Seq data

Compared with five biclustering algorithms (Bimax(18), ISA(41), FABIA(20), Plaid(15), and QUBIC(19), with more details in Table S1 of Supplementary File 1), the performance of QUBIC2 in identifying FGMs was systematically evaluated using four gene expression datasets: a simulated RNA-Seq dataset based on an in-house method (22,846 rows × 100 columns), a bulk RNA-Seq dataset from *Escherichia coli* (*E. coli,* 4,497 rows × 155 columns), a bulk RNA-Seq dataset from TCGA (3,084 rows × 8,555 columns), and a scRNA-Seq dataset from human embryos (3,798 genes × 90 cells). For the identified biclusters from a specific tool, *precision* showcases the fraction of biclusters whose genes are significantly enriched with certain biological pathways (i.e., relevance), and *recall* reflects the fraction of captured known modules/pathways among all known modules in a functional annotation database, e.g., KEGG (42) and RegulonDB (43) (i.e., diversity). The harmonic mean value of precision and recall, referred to as the *F* score, was used as the integrated criteria in performance evaluation.

Evaluation studies usually used default parameters of the to-be-analyzed tools, which were optimized for specific benchmark datasets. However, when applied to datasets coming from a different organism (e.g., *E. coli* vs. human), or be acquired by other technologies (e.g., microarray vs. RNA-Seq), the default parameters often fail to achieve satisfying performance and need further optimization/adjustment. To minimize the biases in performance comparison among multiple tools, for each of the four datasets, we run the six tools under more than 50 parameter combinations by adjusting their critical parameters around default/recommended values (see ***Methods*** and Table S2 in Supplementary File 1). Then the *F* score of identified biclusters under each parameter combination was calculated. In this way, we can test a tool’s robustness and infer how sensitive of its performance is to parameter adjustment, besides the basic performance comparison among different tools.

As showcased in Figure 2, QUBIC2 achieved the highest median *F* scores and the highest *F* scores with the default parameter on all the four datasets, and its *F* scores were significantly higher than the second-best algorithms in all the comparison circumstances (Wilcoxon test *P*-value <0.01). QUBIC2 performed well in both precision and recall, indicating that the identified FGMs are relevant and diverse; and it had relatively small variance, while the performance of some algorithms on certain dataset was very sensitive to parameter change (e.g., FABIA on *E. coli*). Regarding median *F* scores, QUBIC was the second-best algorithm on simulated data, *E. coli* RNA-Seq data, and human scRNA-Seq data, while FABIA was the second-best one for TCGA data. As regards the default settings, QUBIC ranked as the top ones on simulated data and *E. coli* data, and ISA and Plaid had relative higher rank on TCGA data. ISA was generally very stable, and its variances were the smallest on three datasets. As for Bimax, although its recall was relatively low, it was characterized with high precision on the four datasets. It is noteworthy that QUBIC2 is the only program, among all the six biclustering algorithms, which did not encounter a dramatic performance drop on scRNA-Seq data compared to RNA-Seq data, suggesting the unique applicative power of QUBIC2 on FGMs detection from scRNA-Seq data (Figure S1 in Supplementary File1).

**Figure 2.**
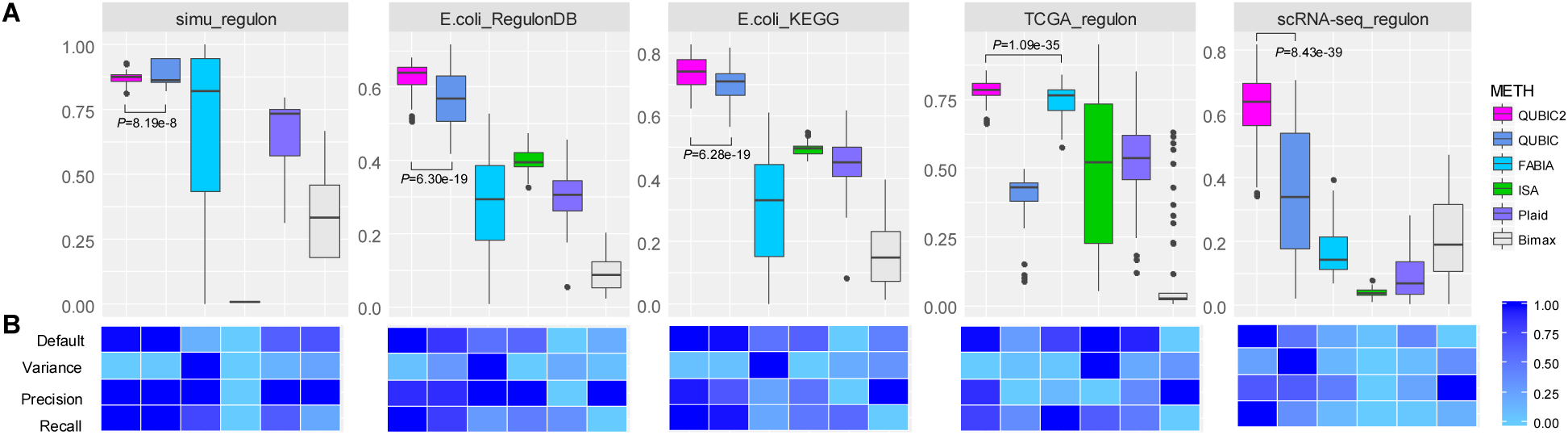
Overall performance comparison between QUBIC2 and five popular biclustering methods based on the agreement between identified biclusters and known modules. **A**. Distribution of F scores on each of the four datasets under multiple runs (n>40). Black line in the box denote median value, whiskers denote 10% and 90% percentiles, while the box denotes 25% and 75% percentiles; **B**. relative performance of six algorithms in terms of F score under default parameters, variance of F scores under multiple sets of parameters, median value for the precision and median value for the recall, respectively (normalized over six algorithms). Note that the variance of F scores depends on the increment of parameters, and therefore only indicative.

Furthermore, the performance of all the biclustering algorithms on *E. coli* data was better than on human data, with the possible reason that *E. coli* data has more completed functional annotation and affects the evaluation of module significance. Therefore, for less annotated organisms, we need a statistical evaluation framework for all the identified biclusters.

### A statistical evaluation framework for identified biclusters

The significances of gene modules from the identified biclusters were usually evaluated by pathway enrichment analysis. However, many organisms (including human) have limited functional annotations supported by experimentally verifications, which makes a systematic evaluation of all identified biclusters non-trivial. To fill this gap, a statistical method was proposed in this study, which can calculate a *P*-value for a bicluster purely based on their size (number of genes and conditions).

Interestingly, we found that there is a strong association between the *P*-values of biclusters calculated via pathway enrichment analysis (named knowledge-based *P*-value) and the corresponding size-based *P*-values. Specifically, spearmen correlation tests were conducted between size-based *P*-values and five groups of knowledge-based *P*-value (see **Methods**). The average spearman correlation coefficients (ρ) were higher than 0.40 (ComTF_ρ =0.48, TF_ρ=0.56, KEGG_ρ =0.42, SEED_ρ=0.43 and ECO_ρ =0.42), and the average *p*-values for the correlation test were smaller than 0.01. As showcased in Figure 3A, all the ρs in the five groups are positive. In addition, ρs related with regulatory pathways (i.e., TF_ ρ and ComTF_ ρ) were generally larger than ρs those related to metabolic pathways (i.e., KEGG_ ρ and SEED_ ρ). This indicated that the size-based *P*-value seemed to be more suitable for the evaluation of biclusters’ regulatory significance. Furthermore, all the corresponding *p*-values were less than 0.05 (Figure 3B), suggesting that the correlations between knowledge-based *P*-values and size-based *P*-values were statistically significant at the 0.05 level. In addition, the parameter *f* which controls the level of overlaps between biclusters had a negative association with ρ (Figure S2 in Supplementary File1), suggesting that the size-based *P*-values would have a stronger association with knowledge-based *P*-values when the overlaps between biclusters are relatively low.

**Figure 3.**
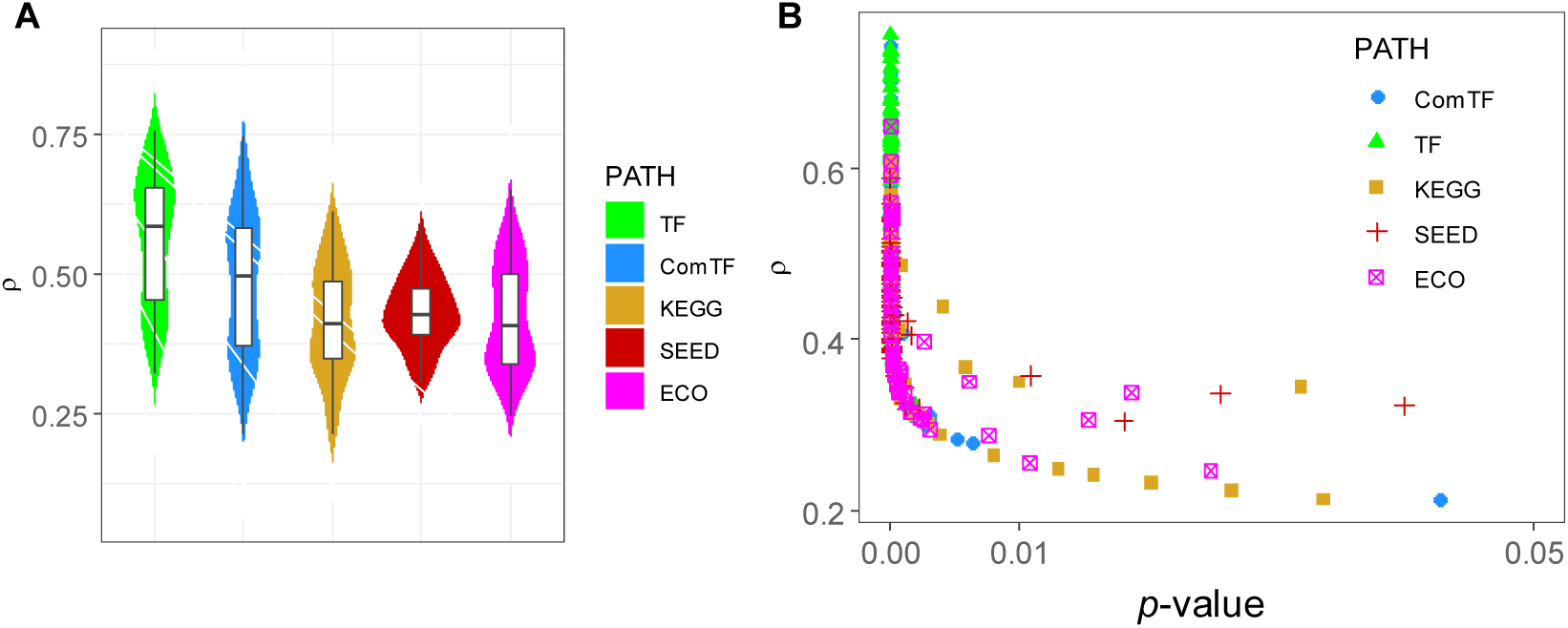
**A**. The distribution of correlation coefficients(ρ) between *P*-value obtained from enrichment analysis and size-based *P*-value. We run QUBIC2 under 63 different parameter settings, and ρ was calculated under each run; **B**. Scatter plot of ρ and p-value. The y-axis denotes ρ, the correlation coefficient for the spearman association test, the x-axis denotes the p-value of the association test. Note that to distinguish, italic lowercase p was used to denote the p-value of the Spearman correlation test, while italic uppercase P was used to denote the significance of biclusters.

### Cell type classification based on scRNA-Seq data

The above sections demonstrated the outstanding performance of QUBIC2 on FGMs identification and its unique feature of statistical evaluation for all the identified biclusters. In this section, we showed the predictive power of biclustering methods on cell types identification from scRNA-Seq data.

We developed a pipeline to group cells into different types with the assumption that two cells belonging to the same bicluster have a higher likelihood of being the same cell type than two randomly selected cells (see **Methods**). Briefly, a biclustering tool was first used to identify biclusters from a scRNA-Seq expression data. Then, a weighted graph *G* = (*C, E*) was constructed to model the relationship between cell pairs, where nodes *C* represent cells, edges *E* connect pairs of cells, and edge weight indicates the number of biclusters that the two corresponding cells appear in simultaneously. Finally, cell types were predicted via the Markov Cluster Algorithm (MCL) clustering on the weighted graph (Figure 4A).

**Figure 4.**
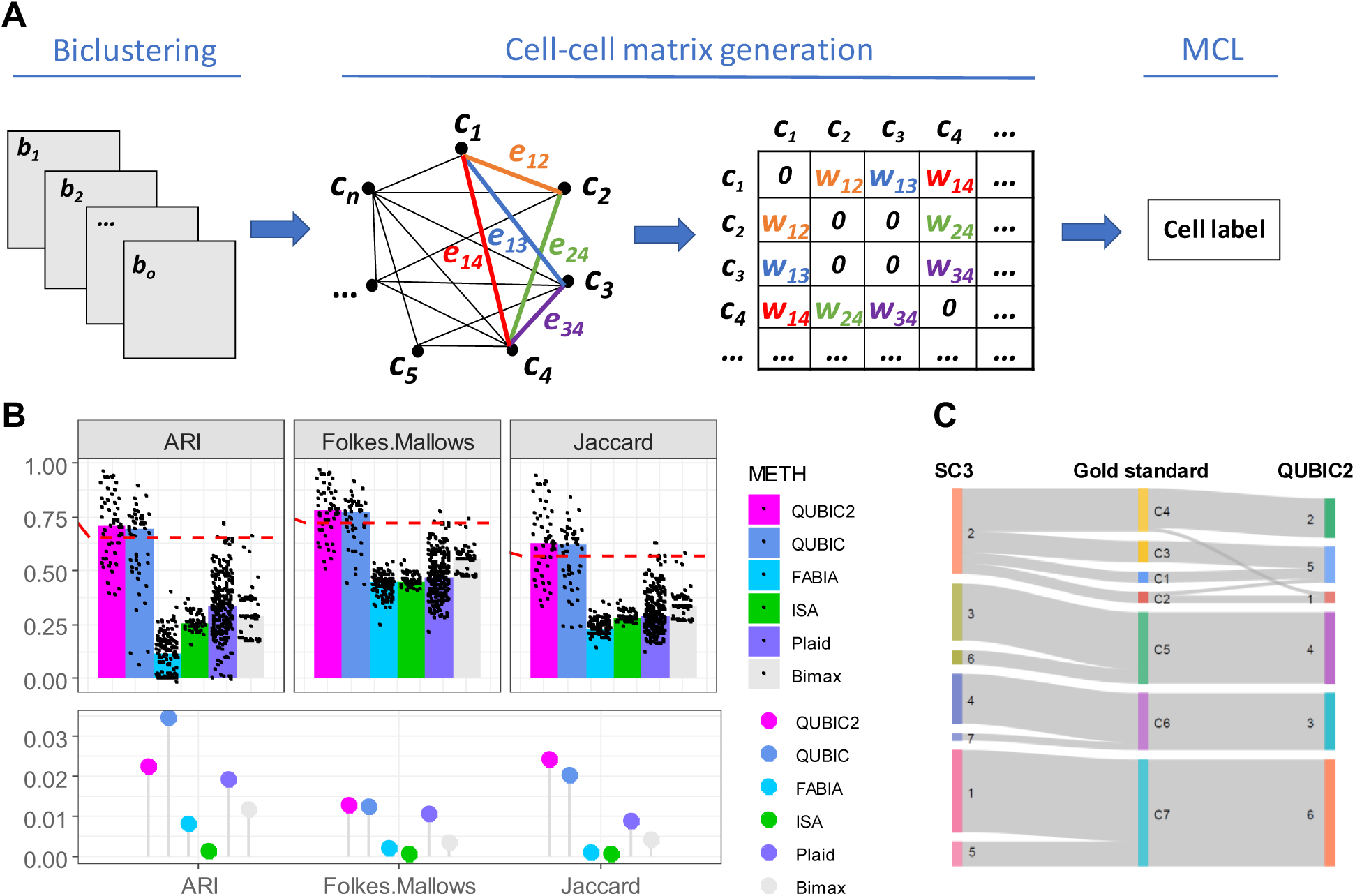
**A**. Computational pipeline for cell type classification. This pipeline consists of three steps: biclustering, generation of weighted cell-cell matrix and clustering using MCL. The input is biclusters and output is cell type labels; **B**. Benchmark of QUBIC2 against five popular biclustering algorithms. Upper layer: each panel shows the similarity between the inferred labels and the reference labels quantified by the four indices, i.e., Adjusted Rand Index (ARI), Folkes Mallows’s index and Jaccard Index, respectively. Each algorithm was applied >40 times to the same dataset to evaluate accuracy and stability. The three indices were calculated for each run of the respective methods (black dots). Bars represent the median of the distribution of black dots. The red dash lines correspond to the benchmark performance of SC3 (ARI: 0.6549, FW: 0.7243, JI: 0.5671). Lower layer: the variance of each tool in terms of three validation criteria;**C.** Sankey diagram comparing the 7 clusters obtained with SC3 (left layer) and 6 clusters obtained with QUBIC2 (right layer). The middle layer corresponds to the seven reference clusters. The widths of the lines linking nodes from two layers correspond to the number of cells they have in common.

For each of the six biclustering methods in Figure 2, we applied this pipeline to a benchmark dataset with 20,214 genes and 90 cells (41), which have been experimentally classified into seven types (46). The Adjusted Rand Index (ARI) was adopted as the evaluation criteria to access the agreement between predicted cell types and these ‘ground truth’ (46). Two more external validation criteria, namely Jaccard Index (JI) and Fowlkes Mallows Index (FW), were also used here aiming to provide a comprehensive evaluation.

As Figure 4B showed, QUBIC2 and QUBIC were the top two biclustering tools, respectively, in terms of median values on the three criteria. Both surpassed the performance of SC3 (41), which was used as the benchmark (median value from 100 runs) and was denoted by the red dash line in each panel of Figure 4B. In addition, ISA always demonstrated the smallest variance across the three validation criteria. The FW values of each tool were more stable than other two values. Figure 4C showcased one cell type classification result obtained by QUBIC2 (parameter *f* was set to 0.85, *c* set to 0.85, *k* set to 13, *o* set to 2000). The result was in good agreement with the reference cell labels and QUBIC2 correctly grouped the three major cell types (8_cell_embryo, Morulae, and late_blastoCyst).

### QUBIC2 inferred the temporal and spatial organization of cells from scRNA-Seq data

When spatial and temporal information is available, scRNA-Seq can reveal more biological insights beyond cell types. In this section, QUBIC2 was applied on two temporal (and) and two spatial scRNA-Seq datasets, respectively, to explore the temporal and spatial organization of cells.

Five biclusters were identified by QUBIC2 from a time series lung scRNA-Seq data (*GSE52583*), which consists of 152 cells collected at E14, E16 and E18, respectively (47). Three of the five biclusters contain time-specific cells. In particular, bicluster BC002 consists of cells exclusively from E14; bicluster BC003 contains cells that only from E16; and bicluster BC004 has cells coming from E18 (Figure 5A). Functional enrichment analyses of the component genes from these three biclusters were carried out based on DAVID (48) and the results showed that genes in BC002 mainly related to cell cycle, cell division, and mitosis; BC003 genes were enriched with ribosome, translation, and structural constituent of ribosome; and spliceosome-related genes were grouped in BC004 (see details in Supplementary File 2).

**Figure 5.**
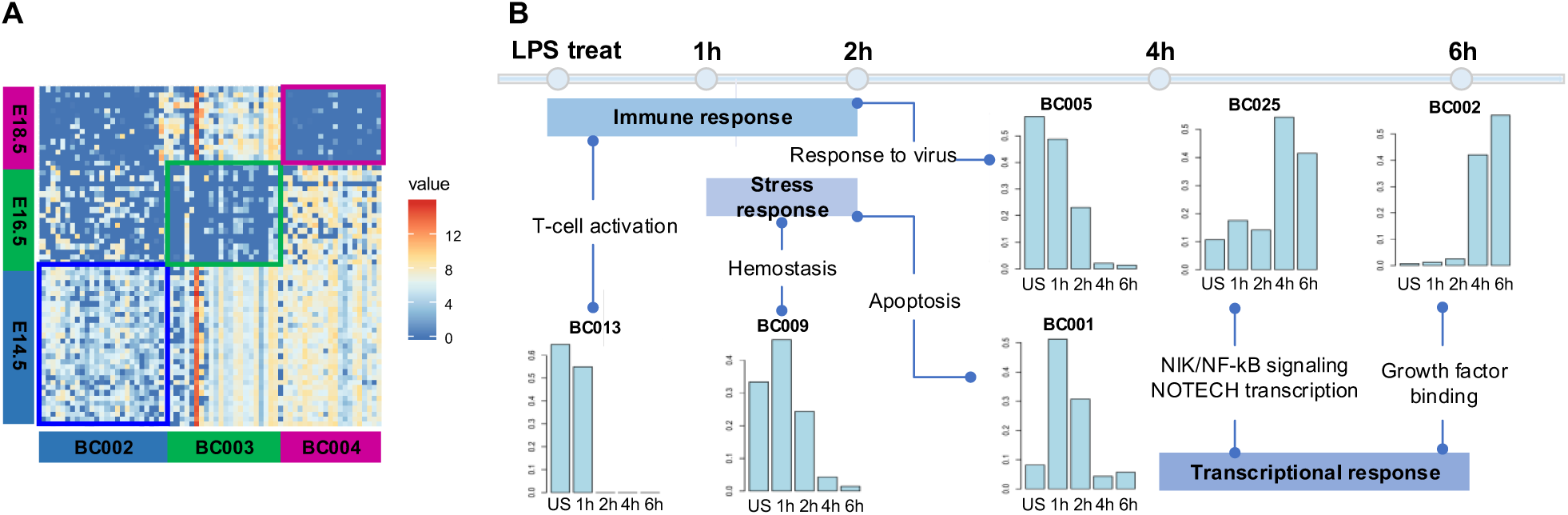
**A**. Visualization of three biclusters (BC002, BC003, and BC004) selected based on the specificity to time point; **B**. Time-dependent distribution of cells in six selected biclusters identified in the LPS data. In each histogram, the five bars from left to right show the proportion of the untreated samples and samples collected at 1h, 2h, 4h and 6h after the LPS treatment.

In addition to identifying biclusters corresponding to specific time point, QUBIC2 can also be used to find biclusters with time-dependent patterns. Here QUBIC2 was used to analyze a scRNA-Seq data with mouse dendritic cells (DCs) collected at 1h, 2h, 4h and 6h after treatment with pathogenic agent lipopolysaccharide (LPS) and untreated controls (*GSE48968*) (49). In total, 51 biclusters were identified in the datasets treated with LPS. For each bicluster, the Fisher exact test was conducted on its constituting samples to assess if significant over-representation by any time points could be found within the bicluster. For those biclusters showing significant association with the time-course, a pathway enrichment analysis was conducted to infer the biological characteristics of the bicluster. In the end, 30 biclusters that are significantly over-represented by one or several consecutive time points were identified in the LPS dataset (α=0.005, *P*<1e-22), and six of them showed distinct time dependence (Figure 5B). Specifically, bicluster BC013 consists of untreated samples and samples collected at 1h, which represents the earliest response to LPS and enriches multiple immune response pathways. Bicluster BC005 consists largely of untreated samples and samples collected at 1h and 2h, which also is enriched with immune response pathways but with more responses to a virus, T cell chemotaxis and so on. BC009 and BC001 are enriched by samples collected at 1h and 2h, covering a wider range of stress-response pathways, suggesting that the activation of stress response pathways and altered metabolisms as secondary responses after the early immune response. BC025 and BC002 consist of samples collected at 4h and 6h, and their genes enrich pathways associated with alterations in cell morphogenesis, migration, cell-cell junction and so on. Overall these observations suggest that our analysis can identify all the major responses to the LPS treatment in a time-dependent manner. Detailed pathways enriched by the six biclusters are given in Figure S3 in Supplementary File 1. The detailed information of these biclusters is given in Supplementary File 3.

Then QUBIC2 was applied to a mouse spatial scRNA-Seq dataset with 280 cells. The cells were classified into five clusters that correspond to five well-defined morphological layers in (50) (Figure 6A). Five biclusters were predicted. Among them, the bicluster BC000 consists of cells mainly from the granular layer; the bicluster BC001 contains cells from the mitral layer and glomerular layer; and the bicluster BC002 contains cells mainly from the olfactory nerve layer (Figure 6B). Functional annotation showed that BC000 mainly enriches plasma membrane, cell membrane, and cell projection; BC001 enriches synapse, neuron projection, and cell projection; and BC002 enriches cell projection (Details in Supplementary File 4).

**Figure 6.**
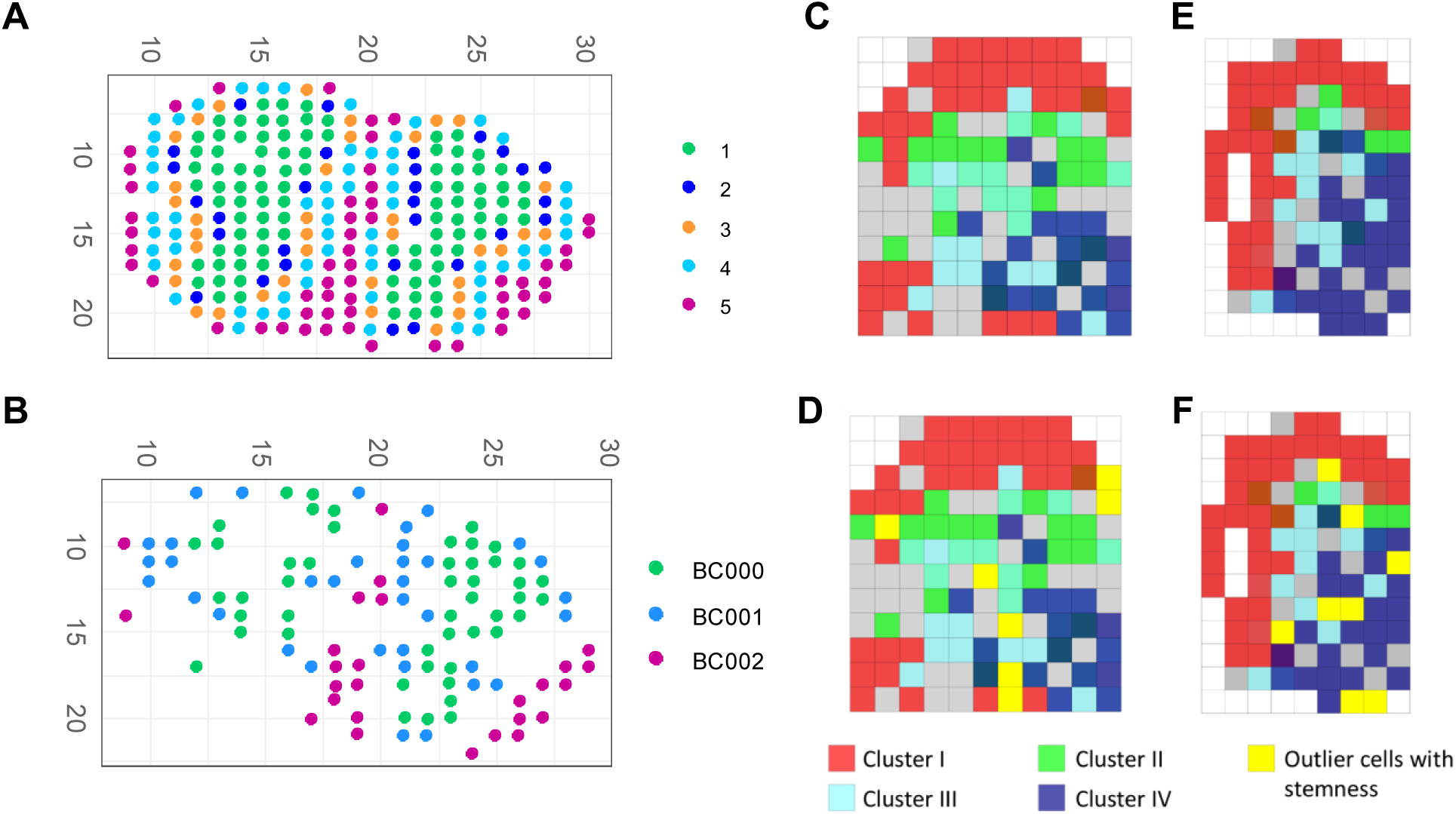
**A**. The coordinates of cells correspond to five morphological layers (1. Granular cell layer; 2. Mitral cell layer; 3. Outer plexiform layer; 4. Glomerular layer; 5. Olfactory nerve layer); **B**. The coordinates of cells from three selected biclusters; **C**. The spatial coordinates of samples in the four biclusters identified in wild-type 1 mouse; Colors red, green, cyan and dark blue represent samples in four different biclusters; **D.** In addition to the coordinates of bicluster samples, the yellow cubes represent significant outlier samples; **E.** The same information as in C except the samples are from wild-type 2 mouse; **F**. The same information as in D except the samples are from wild-type 2 mouse.

Finally, another spatial scRNA-Seq dataset (*GSE60402*) with samples dissected from three mouse medial ganglionic eminence tissues and known spatial coordinates was analyzed. QUBIC2 was applied and 37, 40, and 120 biclusters were identified in the mutant, wild-type 1, and wild-type 2 datasets, respectively (Details in Supplementary File 5). Further investigation on the spatial distribution of cells in each bicluster showed that all the four spatial biclusters with distinct expression patterns by cell cycle, cell morphogenesis, and neuron development genes, as reported in the original study (51), were identified by QUBIC2. It is noteworthy that the outliers with highly expressed stem cell markers tend to be located at the intermediate region between two adjacent (or overlapping) biclusters in the three datasets as shown in Figure 6D and 6F. Our interpretation is that these location-dependent expression patterns may be caused by parallel and independent differentiations from common stem cells.

## METHODS AND MATERIALS

### Data acquisition

A total of four expression datasets were used in the ***Functional gene modules detection from RNA-Seq data*** section, that is, one synthetic RNA-Seq data, one *E. coli* RNA-Seq data and two human datasets (one RNA-Seq and one scRNA-Seq). The synthetic dataset was simulated using our in-house simulation method (details in ***Simulation of co-regulated gene expression data*** section). It contains 22,846 genes and 100 samples. A total of 10 co-regulated modules was embedded in this dataset, covering 2,240 up-regulated genes. The *E. coli* RNA-Seq data consists of 4,497 genes and 155 samples, which was integrated and aggregated by our group. In short, 155 fastq files were downloaded from ftp://ftp.sra.ebi.ac.uk/vol1/fastq/ using the sratoolkit (v2.8.1, https://github.com/ncbi/sra-tools/wiki/Downloads), and they are processed following quality check (FastQC), reads trimming (Btrim), reads mapping (HISAT2) and transcript counting (HTseq). Then, raw read counts were RPKM normalized. The human RNA-Seq data contains 3,084 genes and 8,555 samples, which was obtained from (30). The scRNA-Seq data was downloaded from (13) as an RPKM expression matrix with 20,214 gene and 90 cells and then 3,798 genes were kept for the analysis in this study by removing the genes without annotation.

Multiple sets of known modules/biological pathways were provided or collected to support the enrichment analysis of the above four datasets. For synthetic data, the 10 groups of pre-defined up-regulated genes were used as co-regulated modules. For *E. coli* data, we used five kinds of biological pathways, which are complex regulons and regulons extracted from the RegulonDB database (version 9.4, accessed on 05/08/2017), KEGG pathways collected from the KEGG database (accessed on 08/08/2017), SEED subsystems from the SEED genomic database (accessed on 08/08/2017) (44), and EcoCyc pathways from the EcoCyc database (version 21.1, as of 08/08/2017) (45). Complex regulons were defined as a group of genes that are regulated by the same transcription factor (TF) or the same set of TFs. In total, 457 complex regulons, 204 regulons, 123 KEGG pathways, 316 SEED subsystems, and 424 EcoCyc pathways were retrieved, respectively. For the human TCGA and scRNA-Seq data, we used three sets of modules provided by (30).

One golden-standard scRNA-Seq data (52) was downloaded from https://scrnaseq-public-datasets.s3.amazonaws.com/manual-data/yan/nsmb.2660-S2.csv in the cell type classification section. It consists of 20,214 genes and 90 cells, where the cells were assigned into seven subgroups with the true cell subtypes information provided in (52).

The time series lung scRNA-Seq dataset (GSE52583) with 152 cells and 15,174 genes from was downloaded from (47). The cells were collected at three time points: E14, E16, and E18. Another time series scRNA-Seq data with 527 cells and 13991 genes (GSE48968) was downloaded from the GEO database, in which the RPKM values are available.

The Mouse olfactory bulb spatial transcriptomic data was downloaded from (50), which contains 280 cells and 15,981 genes. Ståhl *et al.* (50) classified the cells into five clusters that correspond

to well-defined morphological layers. The cells use coordinates as IDs, and the cell layers information was manually extracted using the ST viewer (https://github.com/SpatialTranscriptomicsResearch/st_viewer), based on the coordinate information (see Supplementary File 6). The raw reads of mouse spatial scRNA-Seq data GSE60402 was retrieved from the SRA database (53,54), and the RPKM values for it were calculated using software packages TopHat (55) and Cufflink (56). GSE60402 was split into three subsets according to sample information. The detailed information of the selected and split datasets is listed in Table 1.

**Table 1.** Summary of GSE60402

| GEO Accession ID | Data ID | Description | #Cells | #Genes |
| --- | --- | --- | --- | --- |
| GSE60402 | GSE60402-Mutant | From Gfra1 mutant sample | 94 | 11094 |
| GSE60402 | GSE60402-Wildtype1 | From wild type mouse 1 | 124 | 10037 |
| GSE60402 | GSE60402-Wildtype2 | From wild type mouse 2 | 94 | 10714 |

**Table 2.** Main parameters adjusted for each algorithm

| Algorithm | Implementation | Parameters |
| --- | --- | --- |
| Bimax | R package 'biclust' | minr, minc, number |
| ISA | R package 'isa2' | set.seed |
| FABIA | R package 'fabia' | alpha, spl, spz, cyc, p |
| Plaid | R package 'biclust' | row.release, col.release, max.layer |
| QUBIC | R package 'QUBIC' | f, c, k, o |
| QUBIC2 | C++ | f, c, k, o |

### Left Truncated Mixed Gaussian (LTMG) model and qualitative representation

To accurately model the gene expression profile of RNA-Seq and scRNA-Seq data, we specifically developed a mixed Gaussian model with left truncation assumption. Denotes the log transformed FPKM, RPKM or CPM expression values of gene X over *N* conditions as X = {*x*_1_, *⋯x*_*n*_}, we assumed that *x*_*j*_*∈ X* follows a mixture of *k* Gaussian distributions, corresponding to *k* possible TRSs. The density function of *x*_*j*_ is:

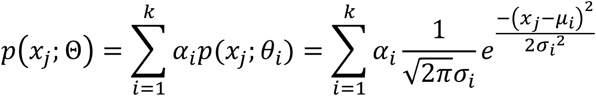

And the density function of X is:

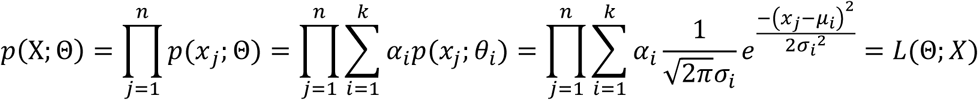

where *α*_*i*_ is the mixing weight, *μ*_*i*_ and *σ*_*i*_ are the mean and standard deviation of *i*^th^ Gaussian distribution, which can be estimated by an EM algorithm with given X:

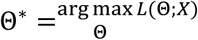

Parameters Θ can be estimated by iteratively running the estimation (E) and maximization (M) steps. In this study, *Z*_*cut*_ is set for each gene as the logarithm of the minimal non-zero RPKM/FPKM/TPM value in the gene’s expression profile. The EM algorithm is conducted for *K* = 1, …, 9 to fit the expression profile of each gene and the *K* that gives the best fit is selected according to the Bayesian Information Criterion (BIC):

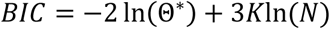

where *K* is the number of TRS, *K* is the number of conditions. *K* that minimizes the BIC will be selected.

Then the original gene expression values will be labeled to the most likely distribution under each cell. In detail, the probability that *x*_*j*_ belongs to distribution *i* is formulated by:

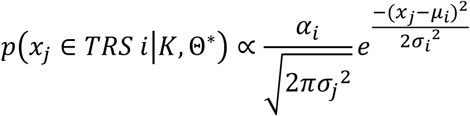

And *x*_*j*_ is labeled by TRS *i* if 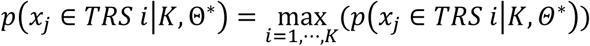. In such a way, a row consisting of discrete values (1,2, ⋯, *K*) for each gene will be generated.

### KL score

A Kullback-Leibler divergence score (**KL** score) is introduced in QUBIC 2 to guide candidate-selection and biclustering optimization. The KL score of a bicluster is defined as:

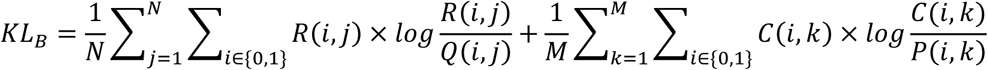

where *N* and *M* are the numbers of rows and columns of a submatrix *B* in *M*_*R*_, respectively. *R*(*i, j*) represents the proportion of element *i* in row *j* of *B, Q*(*i, j*) is the proportion of *i* in the corresponding entire row, *C*(*i, k*) is the proportion of *i* in column *k* of *B*, and *P*(*i, k*) is the proportion of *i* in the entire corresponding column.

Meanwhile, the KL score for a gene quantify the similarity between a candidate gene *j* and a bicluster, which is defined as follows:

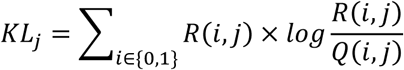

where *R*(*i, j*) represent the proportion of *i* under corresponding columns of the current bicluster.

### Simulation of co-regulated gene expression data

We utilized a single cell RNA-Seq dataset of human melanoma (58) (with 22,846 genes and 4,645 cells) to simulate bulk tissue RNA-Seq data with known co-regulated modules. Specifically, a single cell RNA-Seq pool consists counts data of 4,466 cells of six annotated cell types namely B-, T-, endothelial, fibroblast, macrophage and cancer cells were constructed. The top 1,000 cell type specifically expressed genes of each cell type were identified by using Z score of the mean of each gene’s expression level in each cell type.

For each round of simulation, the number of to be simulated bulk tissue samples and co-regulation modules is first defined. Then the genes of each co-regulation module denoted as *X*, will be specified by randomly selecting *M* genes from the top 1,000 cell type specifically expressed genes of one cell type. A co-regulation strength matrix *P* is then simulated from a bimodal distribution over (0,1), with *P*[*i, k*] denotes the proportion of cells with the transcriptional regulatory signal of co-regulation module *k* in bulk sample *i*. A bulk tissue data is simulated by randomly drawing cells from the cell pool by following a multinomial distribution, with predefined parameters and the total number of cells. For co-regulation module *k* in bulk sample *i*, genes *X* in a proportion *P*[*i, k*] of the selected cells of the cell type corresponds to *k* are perturbed by an X-fold increase of the gene expression. Then the bulk data *i* with simulated co-regulations are formed by summing the perturbed gene expression profile the selected cells and normalized to RPKM expression scale. The Pseudo code of the simulation approach is provided Method S1 in Supplementary File 1.

The rationales of this simulation approach include (1) gene expression level and noise in the bulk data are purely simulated by sum of real single-cell data, without using artificially assigned expressions scale and noise; (2) co-regulation genes are modeled as a specific fold increase of a number of cell-type-specific genes in a particular subset of the cells, which characterizes the heterogeneity of transcriptional regulation among cells in a tissue; (3) multiple co-regulation modules in specific to different cell types can be simultaneously simulated. Hence, we believe the gene expression data simulated by this way can satisfactorily reflect genes co-regulated by a perturbed transcriptional regulation signal in real bulk tissue data.

### Evaluation of the functional modules

The capability of algorithms to recapitulate known functional modules are assessed using precision and recall. First, for each identified bicluster, we use the *P*-value of its most enriched functional class (biological pathway) as the *P*-value of the bicluster. Specifically, the probability of having *x* genes of the same functional class in a bicluster of size *n* from a genome with a total of *N* genes can be computed using the following hypergeometric function(59):

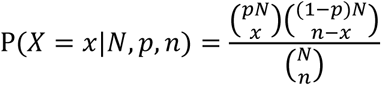

where *p* is the percentage of that pathway among all pathways in the whole genome. The *P*-value of getting such enriched or even more enriched bicluster is calculated as:

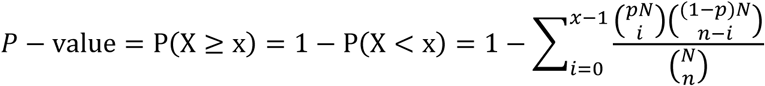

The bicluster is deemed enriched with that function if its p-value is smaller than a specific cutoff (e.g., 0.05).

Given a group of biclusters identified by a tool under a parameter combination, the precision is defined as the fraction of observed biclusters significantly enriched with the one biological pathway/known modules (Benjamini-Hochberg adjusted *p*<0.05)

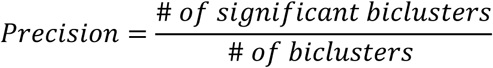

For recall, we compute the fraction of known modules that were rediscovered by the algorithms,

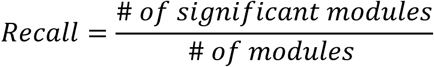

Finally, the harmonic mean of precision and recall were calculated to represent the performance of an algorithm on a given dataset and parameter setting, denoted as *F* score:

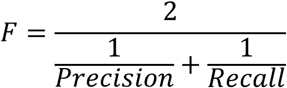

Note that the number of biclusters used to calculate precision and recall may affect the results. To make sure the evaluation is as fair as possible, for each dataset, we select the first 30 biclusters.

### Parameter adjustment of biclustering tools

To assess the robustness of selected algorithms’ performance, each tool is run multiple times by varying parameters that affect the size and number of biclusters. In general, parameters are adjusted around their default or recommended (if available) value. The parameters that varied are listed in Table2, and details about the range and increment of parameters can be found in Supplementary File.

### Spearman correlation test

QUBIC2 was run on the *E. coli* RNA-Seq data in Figure 2 under 63 parameter settings. For each setting, around 100 biclusters were identified. Five sets regulatory or metabolic pathways were extracted from four databases of *E. coli* (RegulonDB, KEGG, SEED (46) and EcoCyc (47)) to support this association study. In specific, for each set of ∼100 biclusters obtained under the same settings, six groups of *P*-values for all these biclusters were calculated, with five knowledge-based groups and one size-based group. Spearman correlation test was conducted to investigate the rank-order correlation among the six groups of *P*-values. Five correlation coefficients (ρ), which demonstrated the extent of correlation between size-based *P*-values and five biological knowledge-based *P*-values, as well as five corresponding *p*-values, were recorded from the test. Note that the *p*-value of correlation test denotes the probability of observing such a correlation or even stronger correlation, under the null hypothesis that no correlation exists. For simplicity, the correlation coefficient between the size-based *P*-value and biological knowledge-based *P*-value was prefixed with the name of a pathway, e.g., TF_ρ and KEGG_ ρ. In the end, a total of 5 × 63 ρ (63 parameter settings, each with five ρs) and a same number of *p*-values were obtained.

### Cell type classification pipeline

By using biclustering, we can group genes and cells simultaneously. However, since biclustering aims to find sets of genes that are co-expressed across a subset of conditions, it is possible that genes may co-expressed across multiple cell types. Therefore, one bicluster may consist of cells from different types, and cells from the same types may appear in different biclusters. In a word, it is not guaranteed that one bicluster corresponds to one cell type. However, it is assumed that two cells from a bicluster are more likely to be of the same subtypes than the two cells that are randomly selected. It is believed that biclusters can capture this feature to some extent. If there are multiple biclusters and when we condense them together, we can distinguish sets of cells belonging to the same type from sets of cells that are grouped by chance.

Based on the above idea, we developed a pipeline to obtain cell type classification based on biclustering results (Figure 4A). First, a biclustering tool was applied to the expression data (rows represent genes and columns represent cells) to identify a set of biclusters. Then a weighted graph *G* = (*C, E*) was constructed to model the relationship between cell pairs among biclusters. A node _*i*_ in *G* represented a cell, and *e*_*i,j*_ represented the edge connecting _*i*_ and _*j*_, where *i* ≠ *j*. We assigned weight *w*_*i,j*_ to *e*_*i,j*_ to represent the number of biclusters that contain both _*i*_ and _*j*_. Intuitively, a higher *w*_*i,j*_ value indicates that *i* and *j* are simultaneously involved in more biclusters, hence, are more likely to be the same cell type than cell pairs with lower weight. A symmetrical cell-cell matrix with diagonal as 0 was then constructed to record *w*_*i,j*_ and Markov Cluster Algorithm (**MCL**) was performed to cluster cells into cell types and produce cell labels. In specific, the MCL clustering was run 100 times by varying inflation factor, resulting 100 cell labels. A binary similarity matrix was constructed for each cell label: if two cells belong to the same cluster, their similarity is 1; otherwise, the similarity is 0. Then a consensus matrix was built by averaging all similarity matrices. The resulting consensus matrix was clustered using hierarchical clustering with complete agglomeration, and the clusters were inferred at the k level of the hierarchy.

### External cluster validity indices

External validation measures the extent to which cluster labels match externally supplied class labels. Generally, they are based on counting the pairs of points on which two classifiers agree/disagree. Denote two partitions of the same data set as R and Q. The reference partition, R, encode the class labels, i.e., it partitions the data into k known classes. Partition Q, in turn, partitions the data into v categories, which is the one to be evaluated.

Adjusted Rand Index (ARI) is defined as

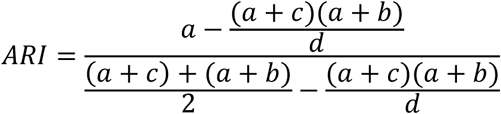

*a*: Number of pairs of data objects belonging to the same class in R and the same cluster in Q.

*b*: Number of pairs of data objects belonging to the same class in R and different clusters in Q.

*c*: Number of pairs of data objects belonging to different classes in R and the same cluster in Q.

*d*: Number of pairs of data objects belonging to different classes in R and different clusters in Q.

Terms *a* and *d* are measures of consistent classifications (agreements), whereas terms b and c are measures of inconsistent classifications (disagreements).

Jaccard Index is defined as:

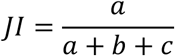

The Jaccard Index can be seen as a proportion of good pairs with respect to the sum of non-neutral (good plus bad) pairs.

Folkes-Mallow’s index is defined as

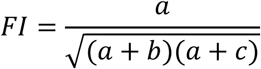

Fowlkes–Mallow’s index can be seen as a non-linear modification of the Jaccard coefficient that also keeps normality.

### Pathway enrichment analysis

Pathway enrichment analysis is conducted and the statistical significance of each enriched pathway is assessed by using a hypergeometric test (statistical significance cutoff = 0.005) against 4,725 curated gene sets in the MsigDB database, which includes 1,330 canonical KEGG, Biocarta and Reactome pathways, and 3,395 gene sets representing expression signatures derived from experiments with genetic and chemical perturbations, together with 6,215 Mouse GO terms each containing at least 5 genes (62,63).

## CONCLUSION

QUBIC2 is a novel biclustering algorithm developed for bulk RNA-Seq and scRNA-Seq data analysis in this study. It has four unique characteristics: (*i*) used a left-truncated mixture model to fit the log-transformed RPKM/CPM/TPM values of each gene and qualitatively represent gene expression; (*ii*) integrates an information-divergence objective function in the biclustering framework; (*iii*) applies a Dual strategy to optimize consistency level of a to-be-identified bicluster; and (*iv*) develops a robust *P*-value framework to evaluate the significance of all the identified biclusters. QUBIC2 proved to have significant advantages in the functional module detection area, outperforming five widely-used biclustering methods based upon our test on four datasets. The proposed *P*-value calculation method based on bicluster size did make sense, which may facilitate the evaluation of all the identified biclusters, especially from less-annotated organisms. The cell type classification pipeline, based on QUBIC2, worked well and outperformed the state-of-the-art performance of SC3. By utilizing time-dependent data, QUBIC2 discovered biclusters specific to time point and identified a cascade of immune responses to the external pathogenic treatment. From the spatial transcriptomic data, QUBIC2 discovered that spatially adjacent single cells may have high co-expression patterns, and particularly, two distinct spatially clustered cells may be derived initially from the same stem cell. We believe that QUBIC2 can serve biologists as a useful tool to extract novel biological insights from large-scale RNA-Seq data (The tutorial for QUBIC2 program is provided in Supplementary File 7).

## DISCUSSION

Single-cell sequencing has enabled new transcriptome-based studies, including the study of distinct responses by different cell types in the same population when encountered by the same stimuli or stresses, and identification of the complex relationships among different cells in complex biological environments such as tissues. However, to fully excavate the potential of scRNA-Seq data, we must overcome several technical challenges.

As sequencing costs decrease, larger scRNA•Seq datasets will become increasingly common; thus, the scalability to large dataset and efficiency of tools will become more and more important. Currently, the discretization and Dual searching functions of QUBIC2 are time consuming on large-scale datasets. Based on our test, it takes 17 minutes to discretize a dataset with 4,297 rows and 466 columns (a desktop with 48.0GB memory, Intel Core i7-6700 and 3.40GHz). Given a dataset with 22,846 genes and 100 conditions, the running time while using Dual strategy are generally 2 minutes longer than that without Dual. The openMP method will be implemented in the EM steps for discretization and more efficient heuristics algorithm will be designed to optimize the dual searching of biclustering.

Another challenge involves the interpretation of time-series and spatial data. For example, in the GSE52583 data, QUBIC2 could only separate cells collected at different time points, yet the further differentiation stage information was not captured. For the mouse olfactory bulb data, QUBIC2 did not separate cells from adjacent layers. To deal with this drawback, we need to combine biclustering with other statistical methods specifically designed for time series and spatial gene expression data.

It is noteworthy that many other kinds of methods can be used for gene expression data analysis. Forty-two module detection tools covering five main approaches were reviewed in (30) and the authors concluded that decomposition methods outperformed all other strategies, including biclustering methods. Meanwhile, they also observed that QUBIC and FABIA had higher performance on human and synthetic data. We compared two top rated decomposition methods and two top clustering methods with QUBIC2 and QUBIC on a human scRNA-Seq data; and the results showed that QUBIC2 surpassed both decomposition and clustering methods (Figure S4 in Supplementary File 1). In the future, we will carry out more comprehensive comparison between QUBIC2 and other decomposition and network-based methods, aiming to give a systematical evaluation of the power of computational techniques on scRNA-seq data.

